# *Trans*-acting role of the leader peptide peTrpL in posttranscriptional regulation of multiresistance

**DOI:** 10.1101/864512

**Authors:** Hendrik Melior, Siqi Li, Konrad U. Förstner, Saina Azarderakhsh, Susanne Barth-Weber, Elena Evguenieva-Hackenberg

## Abstract

Bacterial ribosome-dependent attenuators are widespread posttranscriptional regulators. They harbour small upstream ORFs (uORFs) encoding leader peptides, for which no functions in *trans* are known yet. In the soil-dwelling plant symbiont *Sinorhizobium meliloti*, the tryptophan biosynthesis gene *trpE(G)* is preceded by the uORF *trpL* and is regulated by transcription attenuation according to tryptophan availability. However, *trpLE(G)* transcription is initiated independently of the tryptophan level in the cell, thereby ensuring a largely tryptophan-independent production of the leader peptide peTrpL. We provide evidence that peTrpL plays a role in the differential posttranscriptional regulation of the *smeABR* operon encoding the major multidrug efflux pump SmeAB and the TtgR-type transcription repressor SmeR. We show that peTrpL is involved in a tetracycline-dependent *smeR* mRNA destabilization and forms an antibiotic-dependent ribonucleoprotein (ARNP) complex with *smeR* and its antisense RNA (asRNA). Induction of asRNA transcription, ARNP formation and *smeR* downregulation were promoted by several antibiotics and the flavonoid genistein, and the resistance to these antimicrobial compounds was increased by peTrpL. The role of peTrpL in resistance is conserved in other soil Alphaproteobacteria.

## Introduction

Bacteria resistant to antibacterial drugs (antibiotics) pose an increasing problem for public health and a challenge for development of new antimicrobial strategies or tools [1–4]. In this respect, the discovery of new resistance mechanisms and novel antibiotic interactions is of great interest.

Resistance mechanisms typically comprise changes or protection of the antibiotic target, inactivation of the antibiotic, and prevention of access to target [4,5]. The latter is achieved by increased impermeability for antibiotics, for example by the outer membrane of Gram-negative bacteria, and by active efflux preventing drug accumulation. Chromosomally encoded efflux systems and especially multidrug resistance (MDR) efflux pumps widely contribute to the intrinsic resistance of bacteria towards different antibiotics [4–6], and are of clinical and environmental importance [7]. MDR efflux pumps extrude structurally dissimilar organic molecules such as antibiotics, dyes, solvents, bile or indole [4–7]. Crystallographic analyses of the *Escherichia coli* inner membrane transporter AcrB with ligands revealed the presence of distal and proximal binding pockets, which can accommodate unrelated substrates of different sizes, explaining how an MDR efflux pump provides multiresistance [4,8–10].

In nature, multiresistance is especially important for soil bacteria, since many antibiotic producers are also living in soil [11,12]. Particularly, plant-interacting bacteria have powerful MDR efflux pumps which can also extrude plant antimicrobials such as flavonoids [12–14]. Our model organism, the soil-dwelling plant symbiont *Sinorhizobium meliloti*, harbours genes for 14 MDR efflux pumps [14], but its multiresistance is mainly due to the efflux pump SmeAB, which is also important for nodulation competitiveness. It was shown that deletion of a *smeR* repressor gene, which is located downstream of *smeAB*, increases the resistance to antibiotics, dyes and flavonoids, suggesting that SmeR is the repressor of *smeAB* [14].

Transcription of MDR efflux genes is often induced by binding of the respective antibiotic (efflux pump substrate) to a transcription repressor [4,13,15–17]. The TetR-type repressor TtgR of *Pseudomonas putida*, a bacterium highly resistant to antibiotics, solvents and toxic plant products, is the prototype of a repressor that is capable to bind different classes of antibiotics, which are also substrates of the cognate efflux pump [16]. TtgR crystal structures with ligands showed two distinct and overlapping ligand binding sites, the first broader and hydrophobic, and the second deeper and with polar residues [18]. Thus, in the evolution of Gram-negative bacteria, at least two non-homologous mechanisms for binding of multiple, structurally diverse antibiotics were developed, exemplified by the AcrB-type membrane transporters and the TtgR-type transcription factors. In addition to transcription repression, RND efflux genes are often regulated by global transcription factors like the e.g. MarR-regulated AraC activator [4,19]. However, little is known about the RNA-based regulation of MDR efflux genes.

RNA-based regulation of antibiotic resistance or susceptibility is mediated by *cis*-and *trans*-acting RNAs and RNA-binding proteins. In *cis*, translation inhibition at short upstream ORFs (uORFs) relieves the ribosome-dependent translation or transcription attenuation of downstream resistance genes in Gram-positive bacteria [20–23]. Further, *trans*-acting sRNAs regulate cell wall biosynthesis [24], porine expression [25], and the synthesis of the general stress sigma factor RpoS in enterobacteria, leading to upregulation of the MdtEF efflux pump [26]. Consistently, the RNA chaperone Hfq is important for resistance and is a suitable drug target [27,28]. Furthermore, it is known that *trans*-acting sRNAs and asRNAs are induced upon exposure to antibiotics [29,30]. Thus, antibiotic-induced riboregulators still remain to be uncovered in many bacteria. Other white areas on bacterial chromosome maps are genes encoding small proteins [31,32], which may also be important for resistance.

Small proteins (≤ 50 aa) are poorly characterized or even unknown, because their genes are mostly not annotated [31,32]. However, they can have important functions. Examples are the 26 aa protein SpoVM, which is needed for endospore formation in *Bacillus subtilis* [33], and the 49 aa protein AcrZ, which interacts with AcrB and enhances the export of certain substrates by the *E. coli* AcrAB-TolC pump [34]. The aforementioned uORFs in attenuators [20–23] are common sources of small proteins, the bacterial leader peptides. However, functions in *trans* of bacterial leader peptides are not known yet.

A widespread class of ribosome-dependent transcription attenuators is found upstream of amino acid biosynthesis genes in Gram-negative bacteria [35,36]. The best studied example is the attenuator of the tryptophan (Trp) biosynthesis operon in *E. coli*, which contains the small uORF *trpL* harbouring consecutive Trp codons [37]. Upon *trpL* translation in the nascent RNA, the attenuator can adopt two mutually exclusive structures. When Trp is available, *trpL* translation at the Trp codons is fast, leading to the formation of a transcription terminator and abolishment of the structural genes expression. Upon Trp shortage, the ribosomes transiently stall at the Trp codons, leading to the formation of an antiterminator structure and the structural genes are expressed [36,37].

In *S. meliloti* having three *trp* operons and a *trp* attenuator upstream of *trpE(G)* (Fig. 1A; [38]), the attenuator sRNA rnTrpL, which is generated by transcription attenuation, acts in *trans* to destabilize *trpDC* mRNA [39]. Since the 5′-end of rnTrpL starts with the ATG codon of the *trpL* sORF encoding the leader peptide peTrpL (Fig. 1A and 1B) [38], it may act as a small leaderless mRNA in addition to its role as a riboregulator. In contrast to *E. coli* where transcription of the *trp* genes is repressed under high Trp conditions [36], in *S. meliloti* the *trpLE(G)* operon is not subject of transcription repression [38]. Thus, during exponential growth, *trpLE(G)* is constitutively transcribed and peTrpL is translated at least once per transcription event, independently of Trp availability. This Trp-independent production suggests that peTrpL may have adopted Trp-independent function(s).

**Figure 1.**
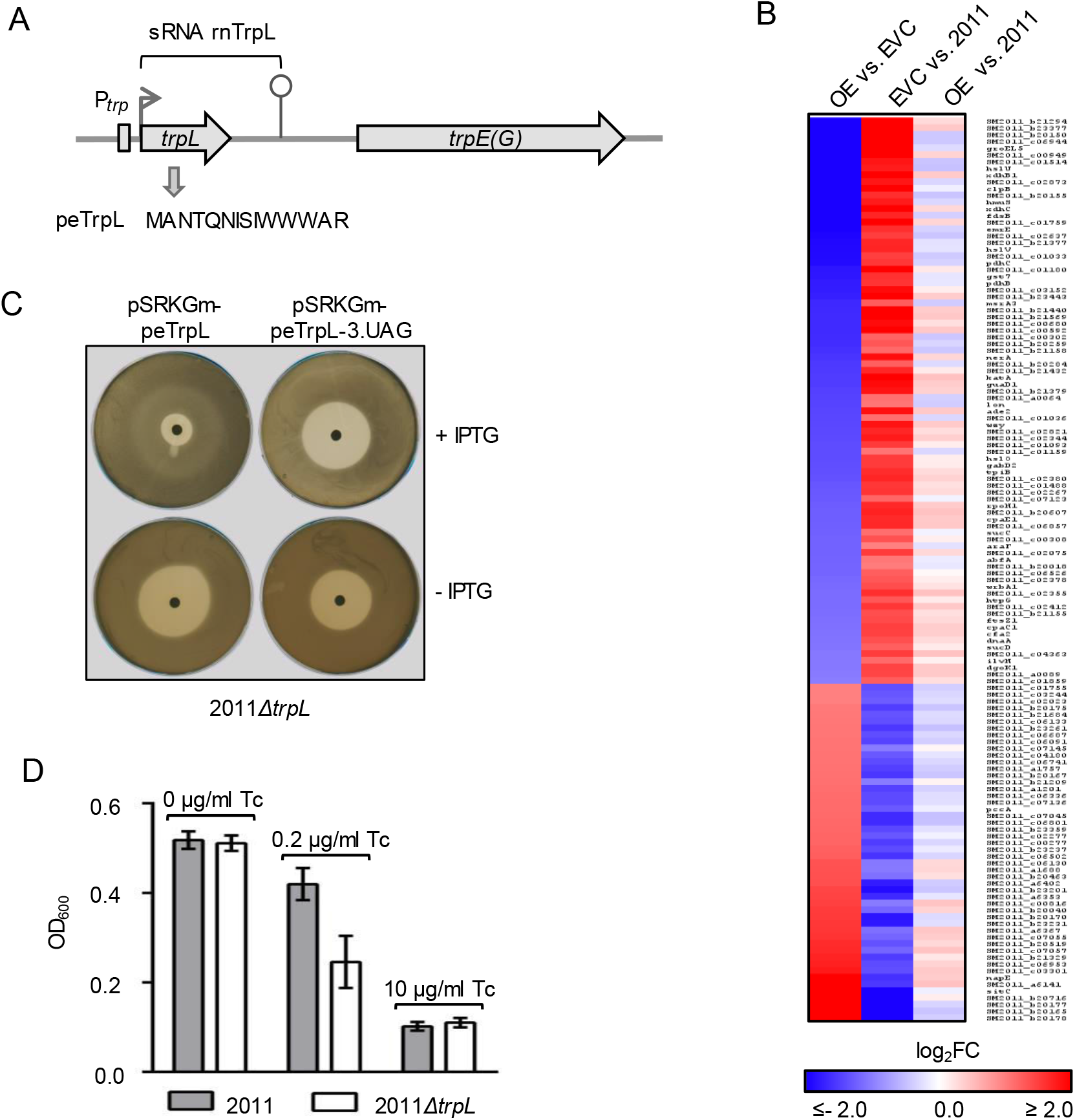
The leader peptide peTrpL increases the resistance to Tc. A Scheme of the *S. meliloti trpLE(G)* locus. The transcription start site and transcription terminator of the *trp* attenuator are depicted by a flexed arrow and a hairpin, respectively (according to [38]). The *trans*-acting products of the *trp* attenuator, the sRNA rnTrpL and the leader peptide peTrpL, are indicated. B Heatmap with RNA-seq data of the following strains: OE, overexpressing strain; EVC, empty vector control; 2011, parental strain. The heatmap shows results for 135 genes with strong differences (log_2_ FC >2.0 or <−2.0) in the comparisons of OE vs. EVC and EVC vs. 2011, which had no or low differences (log_2_ FC >−0.5 or <0.5) in the comparison of OE vs. 2011. C Representative agar plates with zones of growth inhibition by centrally applied Tc. Used strain and plasmids, and presence of IPTG in the agar are indicated. D OD_600 nm_ reached by the indicated strains in microtiter plates, in medium containing the given Tc concentrations. Data from three independent cultures are presented as mean ± s.d.

Here we show that in *S. meliloti*, the leader peptide peTrpL (14 aa) has a role in multiresistance. We found that peTrpL is involved in the antibiotic-dependent destabilization of *smeR* mRNA, which encodes the TtgR-type repressor of the major MDR efflux pump SmeAB. Moreover, we show that peTrpL forms an antibiotic-dependent complex with *smeR* mRNA and an asRNA, which is induced upon antibiotic exposure. This complex is supported by several substrates of the efflux pump SmeAB, including the plant flavonoid genistein. Thus, we uncovered unexpected interactions of antimicrobial compounds with the peptide peTrpL and a target RNAs.

## Results and discussion

### The leader peptide peTrpL increases the resistance to tetracycline

Starting point of this study was our observation that ectopic, constitutive overproduction of the attenuator sRNA rnTrpL (which harbors the ORF *trpL*) from plasmid pRK-rnTrpL apparently counteracts transcriptome-wide effects of tetracycline (Tc) in *S. meliloti*. This observation was based on RNA-seq analysis of the overexpressing strain 2011 (pRK-rnTrpL), the empty vector control (EVC) strain 2011 (pRK4352), and the parental strain 2011. Before harvesting the cells for RNA-seq, strain 2011 was grown in medium with streptomycin (Sm) only, while the strains harboring the Tc-resistance plasmids pRK-rnTrpL and pRK4352 were cultivated in the presence of Sm and Tc. A comparison of the overexpressing strain with the EVC revealed significant changes in the levels of thousands of RNAs (Fig. 1B, Dataset EV1). Surprisingly, when the EVC was compared to strain 2011, inverse changes were observed. Consistently, the transcriptomes of the overexpressing strain and the parental strain 2011 were quite similar (Fig. 1B).

A possible explanation of the differences between the EVC and strain 2011 is a general effect of Tc on mRNA translation [40]. If so, Fig. 1B suggest that the Tc-effect in the overexpressing strain is much lower than in the EVC. Therefore, we hypothesized that overproduction of the sRNA rnTrpL and/or peTrpL peptide encoded by this sRNA may lead to a lower Tc concentration in the overexpressing cells and thus to a higher resistance to Tc. To address this, we used the deletion mutant strain 2011*ΔtrpL* lacking the native rnTrpL RNA being transcribed from the chromosome [39]. To test whether the peTrpL peptide is responsible for the increased resistance, we constructed the gentamycin (Gm) resistance conferring plasmid pSRKGm-peTrpL, which allows for IPTG-inducible peTrpL production. The cloned small *trpL* ORF was mutated to contain synonymous, non-rare codons in order to avoid possible effects by riboregulation or tRNA shortage [39,41]. As a negative control, a plasmid was constructed in which the third codon of the ORF was replaced with a stop codon (pSRKGm-peTrpL-3.UAG). Fig. 1C shows that on plates with centrally applied Tc, the zone of growth inhibition of strain 2011*ΔtrpL* (pSRKGm-peTrpL) was much smaller when IPTG was added to the agar medium. In contrast, the diameter of the bacteria-free halo of the negative control was not decreased on IPTG-containing plates (Fig. 1C). Production of peTrpL also increased the resistance of *S. meliloti* to two natural tetracyclines, chlortetracycline and oxytetracycline (Fig. EV1). We conclude that the leader peptide peTrpL is necessary and sufficient for an increased intrinsic resistance to Tc.

Next we compared the growth of strains 2011 and 2011*ΔtrpL* at different Tc concentrations. Both strains grew similarly in the absence of Tc, and they failed to grow in medium containing 10 μg/ml Tc, which was the half of the concentration used in our selective media. However, in medium supplemented with 0.2 μg/ml Tc, the parental strain 2011 reached a significantly higher OD_600 nm_ compared to the 2011*ΔtrpL* mutant (Fig. 1D). This result provides additional evidence that the native *trpL* is important for the intrinsic resistance of *S. meliloti* to Tc.

### peTrpL is involved in the posttranscriptional regulation the major MDR efflux pump in *S. meliloti*

To address the mechanism by which peTrpL influences resistance, we aimed to coimmunoprecipitate the peptide with its interaction partner(s). For this we used N-terminally tagged 3×FLAG-peTrpL. First we tested whether the tagged peptide is functional. Fig. 2A shows that induced 3×FLAG-peTrpL production increases the Tc resistance of strain 2011 but not of the 2011*Δtrp* mutant, suggesting that the tagged peptide acts in conjunction with the native, wild type peptide. This suggests that peTrpL functions as a dimer or multimer in the cell. Therefore, the coimmunoprecipitation (CoIP) with beads coupled to FLAG-directed antibodies was conducted in the parental background.

**Figure 2.**
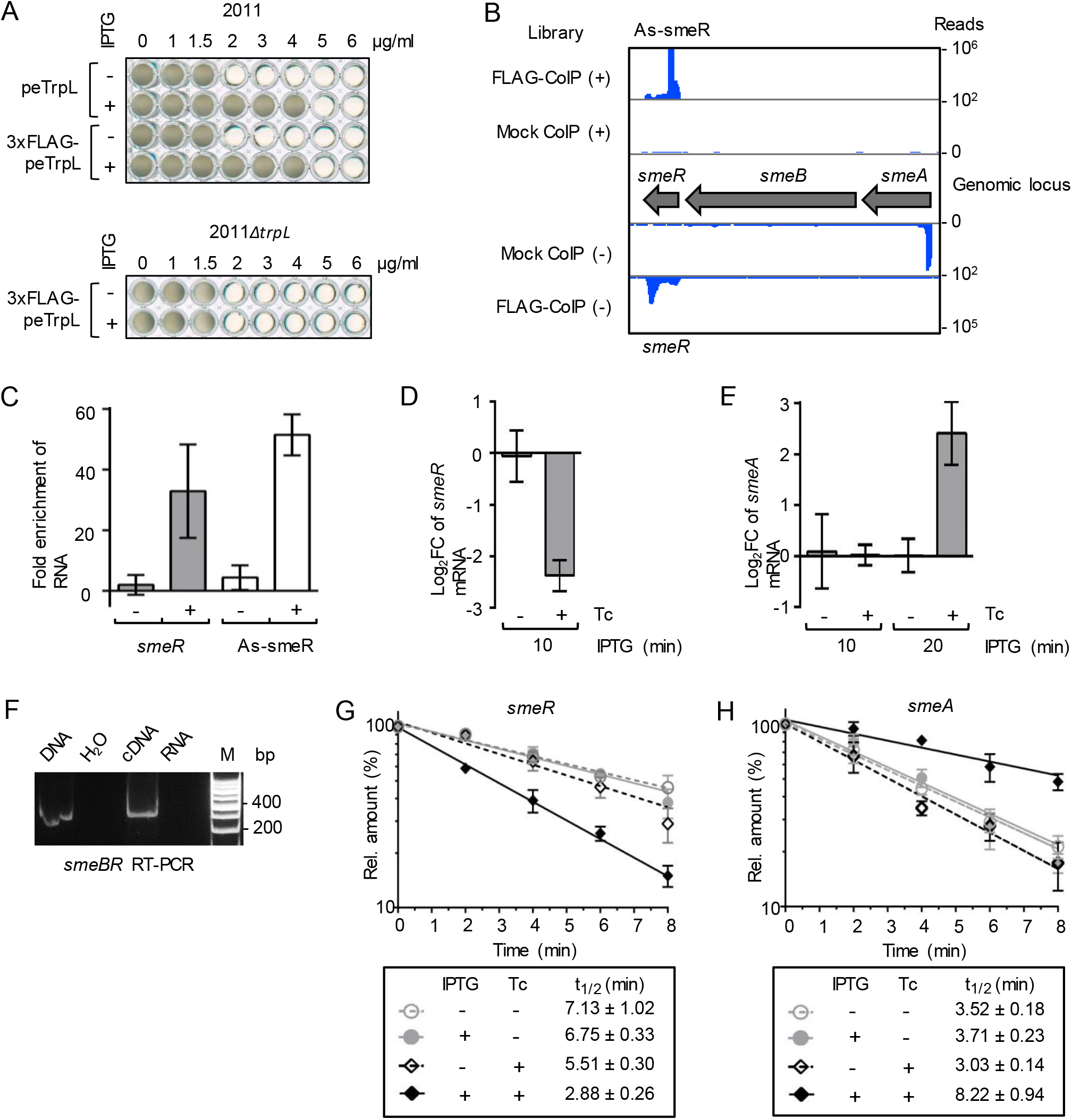
peTrpL posttranscriptionally regulates the *smeABR* operon in a Tc-dependent manner. A Growth of strains 2011 (pSRKGm-peTrpL) and 2011 (pSRKGm-3×FLAG-peTrpL) (upper panels), and 2011*ΔtrpL* (pSRKGm-3×FLAG-peTrpL) (bottom panel) in microtiter plates, in media with increasing Tc concentrations (given above the panels in μg/ml). IPTG presence and peptide products are indicated on the left side. Shown are representative plates. B IGB view of the *smeABR* locus with mapped cRNA reads of the RNA-seq analysis of RNA, which was coimmunoprecipitated with 3×FLAG-peTrpL (FLAG-CoIP). Mock CoIP was conducted with a strain overproducing peTrpL instead of 3×FLAG-peTrpL. Tc was present in the growth medium (20 μg/ml) in the used two-plasmid strains and in the washing buffer (2 μg/ml). C qRT-PCR analysis showing the enrichment of *smeR* and As-smeR RNAs in the FLAG-CoIP samples in comparison to the mock CoIP. Presence of Tc (2 μg/ml) in the washing buffer is indicated. D qRT-PCR analysis of changes in the *smeR* levels 10 min after IPTG addition to two parallel 2011*ΔtrpL* (pSRKGm-peTrpL) cultures. Tc (1.5 μg/ml)) was added together with IPTG to one of the cultures (indicated). E Changes in the *smeA* levels 10 and 20 min after IPTG addition. For other descriptions see D. F RT-PCR analysis with a forward primer located in *smeB* and reverse primer located in *smeR*. The PCR template input is indicated above the panel. G, H Half-lives of *smeR* and *smeA* in strain 2011*ΔtrpL* (pSRKGm-peTrpL) with or without induction of peTrpL production by IPTG. Tc addition (1.5 μg/ml) is indicated. The relative mRNA level values after stop of transcription by rifampicin were determined and plotted against the time. Data information: In (C-E,G,H), data from three independent cultures are presented as mean ± s.d.

**Figure 3.**
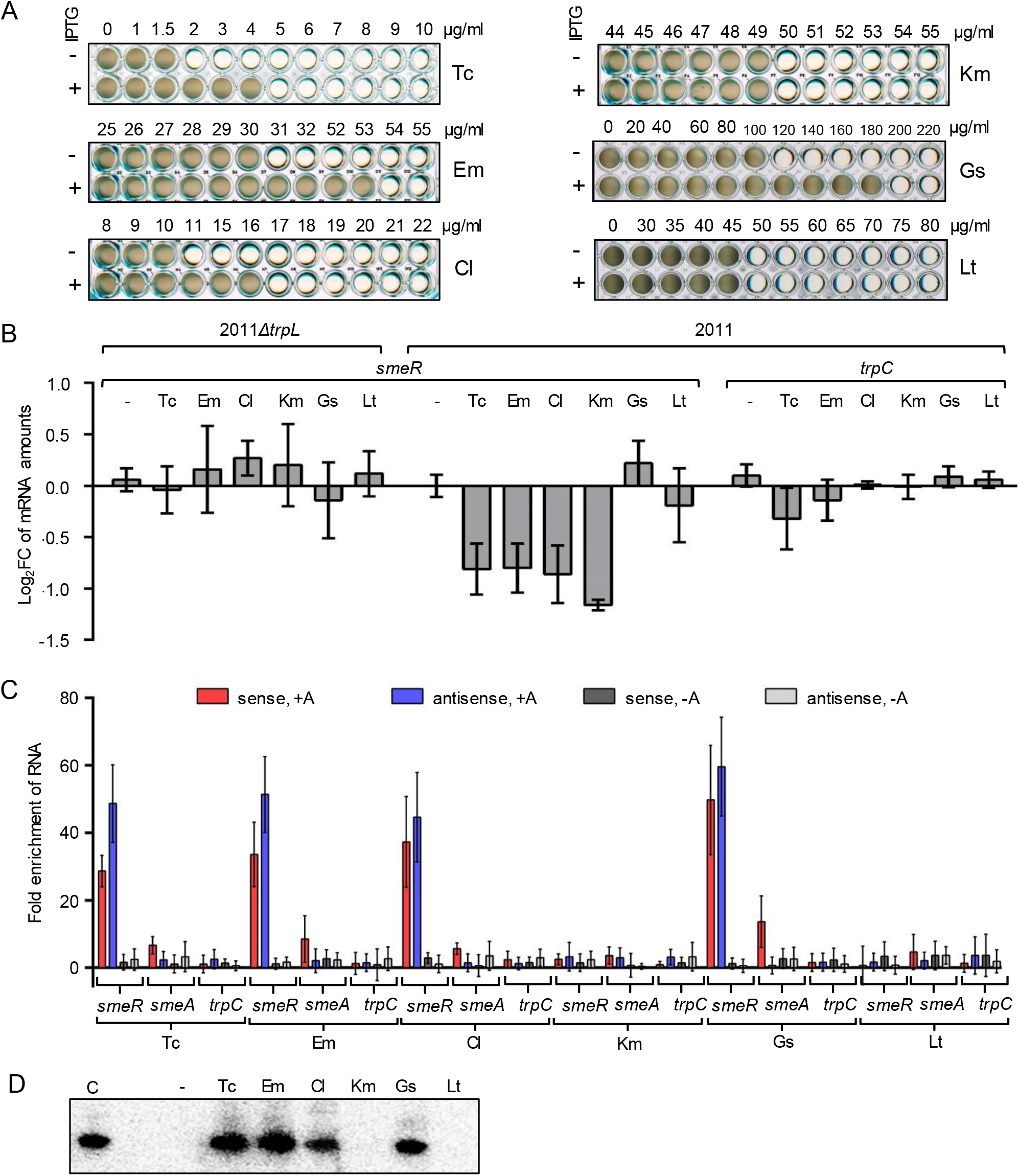
peTrpL increases multiresistance and forms antibiotic-dependent ribonucleoprotein (ARNP) complexes. A Growth of strain 2011*ΔtrpL* (pSRKGm-peTrpL) in microtiter plates. The increasing concentrations of the antibiotics and flavonoids are given above the panels (μg/ml). The antimicrobial compounds are indicated on the right side, IPTG presence and peptide products on the left side of the panels. Shown are representative plates. B qRT-PCR analysis of changes in the *smeR* levels 10 min after addition of the indicated antibiotics and flavonoids (used at subinhibitory concentrations) to cultures of strains 2011*ΔtrpL* and 2011. *trpC*, negative control mRNA. C qRT-PCR analysis of enrichment of the indicated RNAs by FLAG-CoIP in comparison to mock CoIP. Antibiotics and flavonoids (used at subinhibitory concentrations), which were added together with IPTG to cultures of strain 2011 containing either pSRKGm-3×FLAG-peTrpL (FLAG-CoIP) or pSRKGm-peTrpL (mock-CoIP), are indicated below the panel. Presence (+A) or absence (−A) of the antibiotics or flavonoids in the washing buffer are indicated above the panel. D Northern blot analysis of CoIP-RNA from reconstituted ARNP complexes. Used antibiotics and flavonoids are indicated. -, representative negative control of reconstitution, 2 μl ethanol was added to the reconstitution mixture. C, positive control of hybridization, 10 ng of mini-*smeR* was loaded on the gel. Probe directed against mini-*smeR* was used. Shown is a representative hybridization result. Data information: In (B,C), data from three independent cultures are presented as mean ± s.d..

The coimmunoprecipitated RNA (CoIP-RNA) was analyzed by RNA-seq, revealing that *smeR* mRNA encoding the repressor of the major MDR efflux pump SmeAB [14], and a corresponding asRNA As-smeR are potential interaction partners of the peptide (Fig. 2B). This suggests that peTrpL posttranscriptionally regulates *smeR* expression. Surprisingly, we observed that *smeR* and As-smeR were coimmunoprecipitated only if Tc (2 μg/ml, corresponds to the minimal inhibitory concentration) was present in the washing buffer during the CoIP procedure (Fig. 2C). This suggested that Tc is needed for complex formation between peTrpL and the coimmunoprecipitated RNAs, pointing to a direct role of Tc in posttranscriptional regulation.

To test whether peTrpL influences *smeR* expression and whether Tc is needed for this, we analyzed changes in the *smeR* mRNA levels 10 min after IPTG addition to two parallel 2011*ΔtrpL* (pSRKGm-peTrpL) cultures. To one of the cultures, 1.5 μg/ml Tc was added at a subinhibitory concentration together with IPTG. Fig. Fig. 2D shows that the *smeR* level was decreased only if Tc was applied. The gene *smeR* is located downstream of *smeAB* (Fig. 2B) and is cotranscribed with *smeB* (Fig. 2F), We also analyzed the *smeA* level of the tricistronic *smeABR* transcript and found an increase 20 min (but not 10 min) after IPTG and Tc addition (Fig. 2E). Changes in the mRNA levels could be explained by changed mRNA stability. We found that 10 min after induction of peTrpL production, the stability of *smeR* was decreased and that of *smeA* was increased. Importantly, these changes were also dependent on the presence of Tc (Fig. 2G and 2H). While Fig. 2B and 2C suggest that *smeR* mRNA is bound by peTrpL in a Tc-dependent complex leading to its destabilization, the stabilization of *smeA* (probably of *smeAB*) needs further analyses in the future. Altogether Fig. 2 shows that peTrpL and Tc are involved in the differential posttranscriptional regulation of the *smeABR* operon.

Considering the suggested *smeABR* cotranscription upon relieve of repression by SmeR, the need for uncoupling of the expression of *smeAB* and *smeR* at the RNA level is obvious. A Phyre^2^ analysis revealed that SmeR is similar to TtgR (99,9 % confidence; 93 % coverage), the *P. putida* repressor capable to bind different antibiotics [16,42]. This suggests that SmeR allosterically binds Tc and other SmeAB substrates to relieve the repression of *smeABR* transcription [14]. However, antibiotic-induced *smeABR* cotranscription is not favorable for bacterial adaptation under such conditions, when repressor synthesis should be rather avoided in order to ensure increased efflux pump production. The here described destabilization of *smeR* mRNA by a mechanism involving peTrpL and an antibiotic such as Tc serves to downregulate the SmeR synthesis by concomitant SmeAB production (see also Fig. 4E below). The Tc-dependence of this regulation gives a possibility for rapid sensing of the Tc presence in the cell at the level of RNA. When the intracellular Tc concentration is low because enough SmeAB pumps re present in the membrane (and/or because the Tc production in the natural environment was stopped), the *smeR* destabilization by peTrpL and Tc is relieved, the repressor SmeR is synthesized and *smeABR* transcription is repressed again.

**Figure 4.**
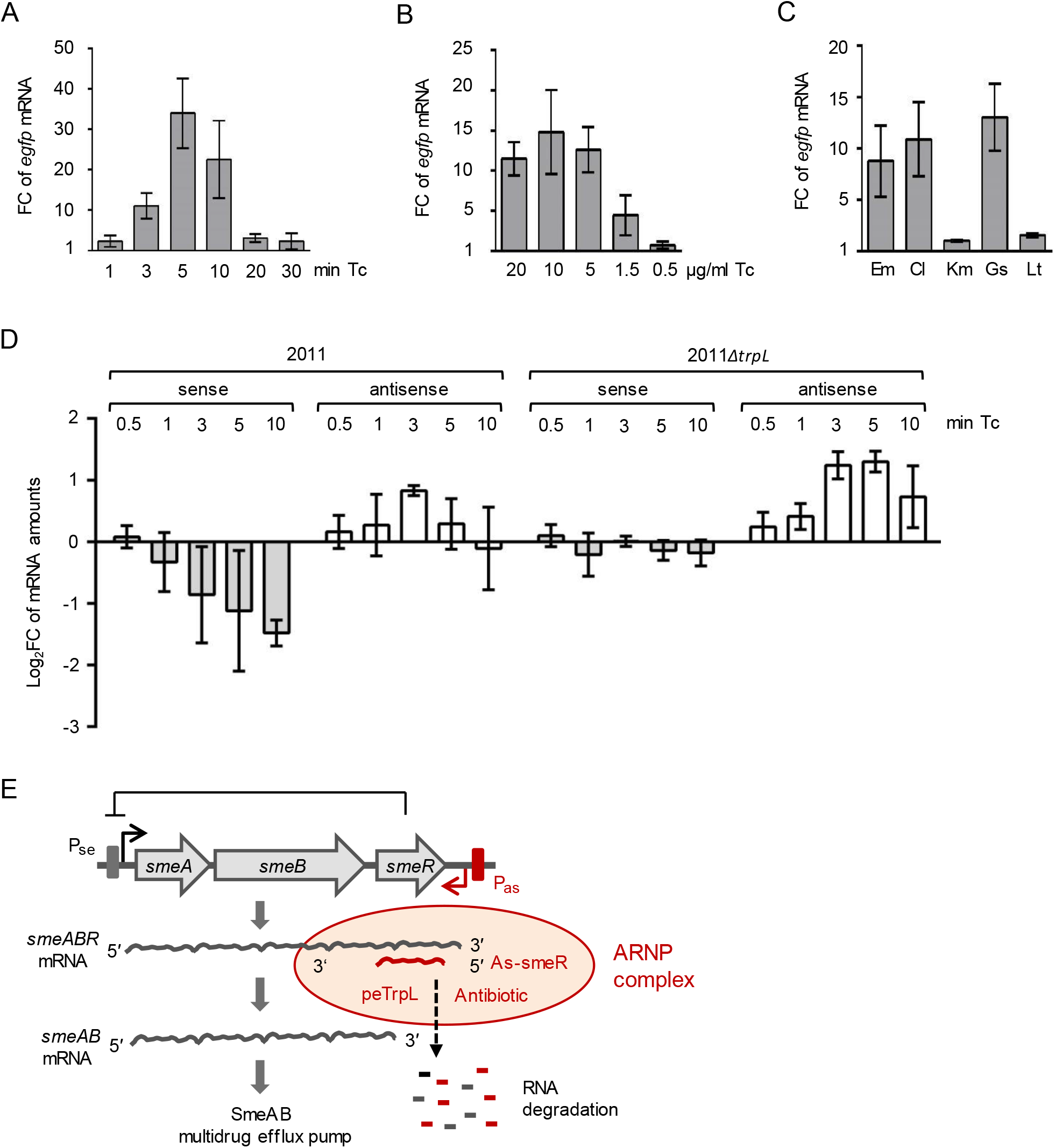
The asRNA As-smeR is induced by substrates of the SmeAB efflux pump. A-C qRT-PCR analysis of reporter *egfp* mRNA reflecting P_as_ promoter activity. A Changes in the *egfp* level upon addition of 20 μg/ml Tc to 2011 (pSUP-PasRegfp) cultures for the indicated time (min Tc). B Changes in the *egfp* level 3 min after addition of Tc to 2011 (pSUP-PasRegfp) cultures. Used Tc concentrations are indicated. C Changes in the *egfp* level 3 min after addition of the indicated antibiotics and flavonoids at subinhibitory concentrations to 2011 (pSUP-PasRegfp) cultures. D qRT-PCR analysis of changes in the levels of *smeR* (sense) and As-smeR (antisense) in the indicated strains upon exposure to Tc (1.5 μg/ml) for the indicated time (min Tc). E Model for the differential posttranscriptional regulation of *smeABR* by peTrpL, antibiotics and the antisense RNA As-smeR. Gene *smeR* encodes the repressor of the *smeABR* operon. Upon exposure to the SmeABR substrates Tc, Em, Cl, or Gs, transcription of *smeABR* and As-smeR from promoters P_se_ and P_as_, respectively, is induced. The peTrpL peptide forms, together with one of the mentioned antibiotics or the flavonoid, an ARNP complex with the *smeR* and As-smeR RNAs, which are then degraded. This leads to *smeR* downregulation and enables the expression of *smeAB* encoding the major MDR efflux pump of *S. meliloti*. Data information: In (A-D), data from three independent cultures are presented as mean ± s.d.

### peTrpL increases multiresistance and forms antibiotic-dependent ribonucleoprotein (ARNP) complexes

According to [14], the antibiotics Tc, erythromycin (Em), chloramphenicol (Cl), and the flavonoid genistein (Gs) are substrates of the MDR efflux pump SmeAB, while kanamycine (Km) and the flavonoid luteolin (Lt) are not. We tested whether peTrpL affects the resistance against these antimicrobial compounds. Fig. 3A shows that peTrpL production increased the resistance of *S. meliloti* to the SmeAB substrates, but not to Km and Lt. Moreover, peTrpL induction increased the cellular efflux (Fig. EV2). These results are in line with a function of peTrpL in the regulation of multiresistance.

The result in Fig. 3A suggests that similar to Tc, the antibiotics Cl and Em, and the flavonoid Gs may participate in the peTrpL-dependent *smeR* downregulation. To test this, we exposed strain 2011 and the 2011*ΔtrpL* mutant to subinhibitory concentrations of antibiotics and flavonoids for 10 min, and analyzed changes in the levels of *smeR* mRNA. Indeed, exposure of strain 2011 to the SmeAB substrates Tc, Em, Cl, and Gs led to a *smeR* decrease, while Km and Lt had no effect (Fig. 3B). The mRNA *trpC* was essentially not affected, showing the specificity of *smeR* downregulation upon exposure to SmeAB substrates. Importantly, the *smeR* decrease was not observed in the 2011*ΔtrpL* mutant, confirming the involvement of peTrpL in this regulation (Fig. 3B).

Next we used CoIP with 3×FLAG-peTrpL to test whether all analyzed SmeAB substrates support a complex of the peptide with *smeR* mRNA and its asRNA. As controls, *smeA* and *trpC* were analyzed. To cultures of strain 2011 (pSRKGm-3×FLAG-peTrpL), IPTG was added together with one of the antibiotics at a subinhibitory concentration. After 10 min, FLAG-CoIP was conducted. The washing procedure was performed either with the respective antibiotic in the buffer, or without an antibiotic, and CoIP-RNA was analyzed. Fig. 3C shows that, provided the the respective antibiotic or Gs was present in the washing buffer*, smeR* mRNA and its asRNA were strongly enriched by the CoIP from cultures exposed to Tc, Em, Cl, or Gs. The specificity of this CoIP is clearly shown by the failure to enrich *trpC*. In comparison to *smeR*, the *smeA* mRNA was enriched only very weakly in the presence of Tc, Em, Cl, or Gs, probably because of the induced *smeABR* cotranscription. Km and Lt did not lead to CoIP of the analyzed RNAs (Fig. 3C). These results suggest the existence of antibiotic-dependent ribonucleoprotein (ARNP) complexes comprising peTrpL, *smeR* mRNA, the asRNA As-smeR, and one of the antibiotics Tc, Em, Cl or the flavonoid Gs.

To confirm ARNP complex formation *in vitro*, we performed reconstitution using synthetic components. Fig. 2B (see above) shows a high, 70 nt peak in the RNA-seq data of the coimmunoprecipitated asRNA As-smeR. We reasoned that this peak may correspond to the binding site of peTrpL to As-smeR and synthesized a corresponding *in vitro* transcript, which we named mini-As-smeR. A complementary *in vitro* transcript was also synthesized, which corresponds to a part of *smeR* mRNA, and was named mini-*smeR*. Both transcripts were mixed with synthetic wild type peTrpL and 3×FLAG-peTrpL and each of the above mentioned antibiotics and flavonoids was added separately. To test for complex formation, FLAG-CoIP followed by Northern blot analysis of mini-*smeR* was performed. According to Fig. 3D, mini-*smeR* was coimmunoprecipitated only if Tc, Em, Cl, or Gs was added. This result shows that these compounds (but not Km and Lt) support ARNP complex formation. It is in perfect agreement with the CoIP data (Fig. 3C) and with the *in vivo* results (Fig. 3A and 3B), strongly suggesting that the SmeAB substrates Tc, Em, Cl and Gs directly participate in the posttranscriptional downregulation of *smeR* by the peTrpL peptide.

### The antisense RNA As-smeR is induced by substrates of the SmeAB efflux pump

The asRNA As-smeR is a part of the ARNP complex that, according to the above data, is formed upon exposure to certain antibiotics or flavonoids. This asRNA was not detected in a previous high-throughput study [43], suggesting that it is induced under specific conditions. To test whether an antibiotic-inducible, antisense promoter (P_as_) is present downstream of *smeR*, plasmid pSUP-PasRegfp, which harbors a transcriptional fusion of *egfp* to the putative P_as_ was integrated in the chromosome of strain 2011. Upon exposure to Tc, the level of the reporter *egfp* mRNA was transiently increased (Fig. 4A), with a significant increase already at 3 min, peak at 5 min, and almost no increase at 20 min exposure time, probably because at the last time point Tc was already pumped out from the cells. Fig. 4B shows the P_as_ induction upon 3 min exposure to different Tc concentrations, including the subinhibitory concentration of 1.5 μg/ml, which was used in most experiments. Further, Em, Cl and Gs (but not Km or Lt) also induced transcription from P_as_ (Fig. 4C).

Next we analyzed kinetics of *smeR* and As-smeR changes in strains 2011 and 2011*ΔtrpL* upon exposure to Tc. In strain 2011, a decrease of *smeR* was detectable already at the time points 3 and 5 min, but the determined values varied, while at 10 min a significant decrease was observed (Fig. 4D). In contrast, a significant increase in the As-smeR mRNA level was transiently detectable after 3 min (but not 10 min) exposure. However, according to Fig. 3C, As-smeR was coimmunoprecipitated with 3×FLAG-peTrpL after 10 min exposure to Tc. We suggest that at 10 min exposure of strain 2011, the As-smeR RNA is a subject of strong co-degradation with *smeR* mRNA, and therefore an increase in the As-smeR level is hardly detectable (Fig. 4D). In strain 2011*ΔtrpL,* the asRNA induction was detectable even 10 min after Tc exposure (Fig. 4D). This longer detection could be attributed to lack of co-degradation with *smeR* (as expected, *smeR* was not downregulated in this strain), but also to less efficient Tc efflux due to lack of the peTrpL-dependent, differential *smeABR* regulation, which, according to our data, is important for the intrinsic Tc resistance.

Altogether, the presented results show a transient production of the asRNA As-smeR upon exposure to SmeAB substrates, and suggest that this asRNA participates together with the inducing antibiotic and peTrpL in the *smeR* downregulation (Fig. 4E).

### Conservation of the peTrpL role in resistance

To test whether the role of peTrpL in resistance is conserved in other bacteria, we used *Agrobacterium tumefaciens* (which, together with *S. meliloti*, belongs to the *Rhizobiaceae*), and the more distantly related *Bradyrhizobium japonicum* (a *Bradyrhizobiaceae* member). In both species, the mRNA levels of their *sme*R homologs were specifically decreased upon overproduction of the corresponding leader peptides Atu-peTrpL and Bja-peTrpL (Fig. 5A). Furthermore, the Tc resistance of both *A. tumefaciens* and *B. japonicum* was increased upon overproduction of their peTrpL homologs (Fig. 5B and 5C). However, production of Atu-peTrpL and Bja-peTrpL in *S. meliloti* did not increase multiresistance (Fig. EV3), pointing to a specificity of interaction between the leader peptides with their targets. Thus, the role of peTrpL in multiresistance is conserved in other soil Alphaproteobacteria.

**Figure 5.**
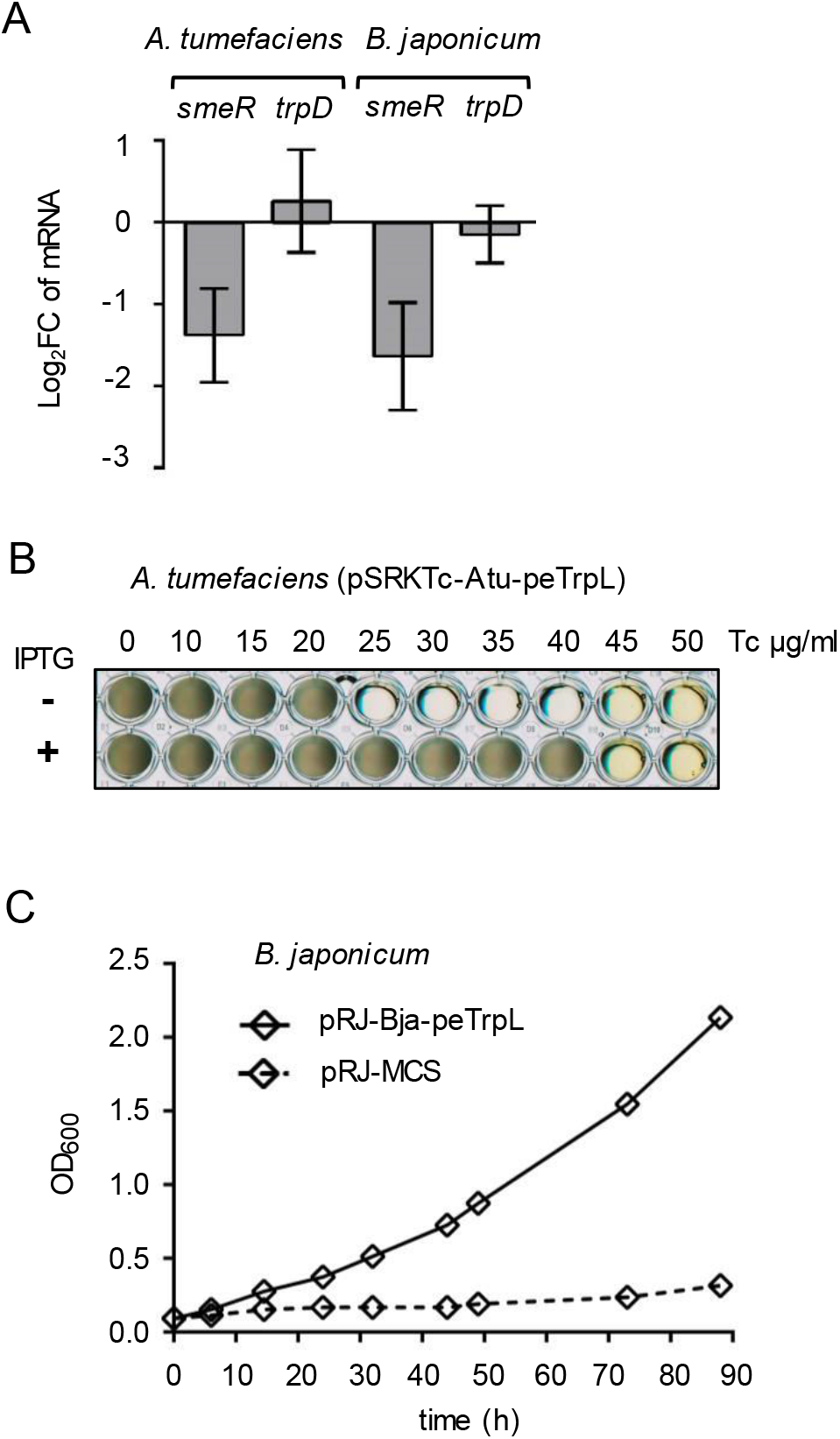
Conservation of peTrpL function in resistance. A qRT-PCR analysis of the expression of *smeR* homologs and *trpD* upon overproduction of the respective peTrpL homologs in *A. tumefaciens* and *B. japonicum*. Data from three independent cultures are presented as mean ± s.d. B Growth of the indicated *A. tumefaciens* strain in microtiter plates. Presence of PTG in the media and the used Tc concentrations are indicated. C Growth curves of *B. japonicum* containing the indicated plasmids (pRJ-MCS, empty vector). Medium supplemented with 100 μg/ml Tc was used. Data from three independent cultures are presented as mean ± s.d. The s.d. were smaller than the symbols in the graph.

In summary, our work establishes a role for the leader peptide peTrpL in *trans*. We provide evidence for a peTrpL function in differential posttranscriptional regulation of the MDR efflux operon *smeABR*. Surprisingly we found that peTrpL forms antibiotic- and flavonoid-supported ribonucleoprotein complexes (ARNPs) with the traget mRNA *smeR* and an asRNA. This suggests that in addition to MDR efflux pumps and multidrug-binding TtgR-type repressors, bacteria developed a third multiresistance-related mechanism for binding of structurally different organic molecules. We propose that the ARNP formation is a novel mechanism for direct utilization of antimicrobial compounds in posttranscriptional regulation of multiresistance. The presented data underline the importance of soil bacteria as a reservoir of new resistance mechanisms [6,7,11]. The conservation of the role of peTrpL in several Alphaproteobacteria encourages future investigations of the ARNP prevalence and the potential of their peptide and RNA components as drug targets.

## Materials and Methods

### Cultivation of bacteria and exposure to antimicrobial compounds

Strains used in this work are listed in Appendix Table S1. *Sinorhizobium (Ensifer) meliloti* 2011 [44,45]; *Agrobacterium tumefaciens* (*A. fabrum*) NTL4 (pZLR4) [46,47] and their derivatives were cultivated in TY medium [48] at 30 °C, *Bradyrhizobium japonicum* (*B. diazoefficiens*) 110*spc*4 [49,50] in PSY medium [51] at 30 °C, and *E. coli* in LB medium at 37 °C [52]. Liquid cultures of Alphaproteobacteria were cultivated semiaerobically (30 ml medium in a 50 ml Erlenmayer flask at 140 r. p. m.) to an OD_600_ of 0.5, and then processed further. For growth experiments in 96-well microtiter plates, 300 μl culture (diluted to an OD_600_ of 0.1) per well was used. Plates were incubated on the shaker (140 r. p. m.) at 30 °C for 60 h (till the cultures entered the stationary phase). Three independent experiments revealed similar results.

Following selective antibiotics concentrations were used when resistance was encoded on a plasmid or the chromosome: tetracycline (Tc, 20 μg/ml for *S. meliloti* and *A. tumefaciens*; *B. japonicum* was cultivated with 25 μg/ml Tc in liquid and 50 μg/ml Tc on plates), gentamycin (Gm, 10 μg/ml in liquid cultures and 20 μg/ml in plates), streptomycin (Sm, 250 μg/ml), spectinomycin (Sp, 100 μg/ml). Following subinhibitory concentrations of antibiotics and flavonoids were used: 1.5 μg/ml Tc, 27 μg/ml Em, 9 μg/ml Cl, 45 μg/ml Km, 90 μg/ml Gs, and 45 μg/ml Lt. Other concentrations used are given in the figures and their legends. The time of exposure to antibiotics and flavonoids is indicated. Tc (tetracycline hydrochloride) and Km (kanamycin sulfate) were purchased from Roth (Karlsruhe, Germany). The other antibiotics including chlortetracycline hydrochloride and oxytetracyline hydrochloride, and the flavonoids were purchased from Sigma Aldrich. IPTG was added to a final concentration of 1 mM.

For the zone of growth inhibition tests, strains 2011*ΔtrpL* (pSRKGm-peTrpL) and 2011*ΔtrpL* (pSRKGm-peTrpL-3.UAG) were used. 15 ml bottom TY agar was overlayed with 10 ml TY top agar mixed with 1 ml *S. meliloti* culture (OD_600nm_ of 0.5). The bottom and the top agar were supplemented with 20 μg/ml Gm. After solidification of the top agar, a Whatman paper disk was placed in the middle of the plate and 5 μl Tc solution (10 μg/μl in 70% ethanol) were applied onto the disk. Plates were incubated over night at 30 °C, before measuring the diameter of the zone of growth inhibition. Three independent experiments revealed similar results.

### Cloning, plasmid characteristics and conjugation

Cloning in *E. coli* was performed by standard procedures [52]. FastDigest restriction enzymes and Phusion polymerase (Thermo Fisher Scientific) were used. PCR amplicons were first cloned in pJet1.2/blunt (CloneJet PCR Cloning Kit; Thermo Fisher Scientific) and then subcloned into conjugative plasmid. For cloning of the *trpL* ORFs with codons exchanged for synonymous codons, or with mutated codons, complementary oligonucleotides were annealed and cloned directly into the desired conjugative plasmids. For *S. meliloti* and *A. tumefaciens*, following conjugative plasmids were used: pSRKGm and pSRKTc (autonomously replicating; allow for IPTG-inducible expression and confer resistance to Gm and Tc, respectively, [53]) or pSUP202pol4 (suicide plasmid, allows for construction of chromosome integrating plasmids; confers resistance to Tc, [54]. Further, the empty vector pRK4352 which confers resistance to Tc [55], was also used, particularly in two-plasmid strains containing in addition pSRKGm-constructs, when it was desired to cultivate bacteria in medium containing 20 μg/ml Tc. Its derivative pRK-rnTrpL was used for constitutive overproduction of the attenuator sRNA rnTrpL and the leader peptide peTrpL encoded by this sRNA [56, 39]. For *B. japonicum*, the chromosome integrating plasmid pRJPaph-MCS was used [57], which confers resistance to Tc. Insert-containing plasmids were analyzed by Sanger sequencing (sequencing service by Microsynth Seqlab, Göttingen, Germany) with plasmid-specific primers. The used oligonucleotides (primers) were synthesized by Microsynth, Balgach (Switzerland). They are listed in Appendix Table S2. The used plasmids are listed in Appendix Table S1.

For IPTG-inducible peptide production in *S. meliloti* or *A. tumefaciens*, small, leader peptide encoding ORFs were cloned in frame using the NdeI restriction site of the plasmids pSRKGm and pSRKTc, which provides an ATG codon [53]. Plasmids pSRKGm-peTrpL and pSRKGm-peTrpL contain an ORF encoding the *S. meliloti* peTrpL peptide. In plasmid pSRKGm-peTrpL-3.UAG, the third codon of this ORF was exchanged for a stop codon, while plasmid pSRKGm-3xFLAG-peTrpL harbors N-terminally fused codons for a triple FLAG-tag. Plasmids pSRKTc-Atu-peTrpL and pSRKTc-Bja-peTrpL contain ORFs encoding the *A. tumefaciens* and *B. japonicum* peTrpL homologs Atu-peTrpL and Bja-peTrpL, respectively. For peptide overproduction in *B. japonicum* and because of the lack of a suitable inducible plasmid for this organism, the ORF encoding the *B. japonicum* peTrpL homolog (Bja-peTrpL) was cloned in pRJPaph-MCS downstream of a constitutive promoter [57]. Compared to the respective chromosomal copy, the recombinant, leader peptide encoding ORFs contained several synonymous nucleotide substitutions (to avoid RNA-based effects [39]), but lacked rare codons to avoid toxicity [41].

To construct plasmid pSUP-PasRegfp, which harbors a transcriptional fusion of *egfp* to the promoter P_as_, 290 bp upstream of the putative TSS of the asRNA As-smeR (according to the RNA-seq data, see also Fig. 2B) was amplified; the first two nucleotides downstream of the putative TSS were included in the reverse primer (Appendix Table S2), which in addition contained the sequence of a typical bacterial ribosome-binding site and the 5′-sequence of *egfp*. The *egfp* sequence of plasmid pLK64 [58] was also amplified, and the two amplicons were used for overlapping PCR. The resulting PCR product was cloned into plasmid pSUP202pol4-exoP, which is a pSUP202pol4 [54] derivative containing 300 nt of the 3′ *exoP* region as a suitable chromosomal integration site [59].

Plasmids were transferred from *E. coli* to *S. meliloti*, *A. tumefaciens* or *B. japonicum* by diparental conjugation with *E. coli* S17-1 as the donor [60]. Bacteria were mixed, washed in saline and spotted onto a sterile membrane filter, which was placed onto a TY plate without antibiotics. After incubation for at least 4 h (for *S. meliloti* and *A. tumefaciens*) or 3 days (for *B. japonicum*) at 30 °C, serial dilutions were spread on agar plates with selective antibiotics.

### Nile red efflux assay

The efflux assay was performed essentially as described [61]. Cultures of strains 2011 (pSRKGm-peTrpL, pRK4352) and the EVC 2011 (pSRKGm, pRK4352) were cultivated in medium with Gm and Tc. Cultures with and without IPTG, which induces peTrpL production, were grown in parallel. Pellets from 20 ml culture were washed in 20 mM potassium phosphate buffer (pH 7.0) containing 1 mM MgCl_2_ (PPB-Mg), and resuspended in PPB-Mg adjusting the OD_600nm_ to 1.0. The cell suspension was incubated for 15 min at room temperature. 2 ml aliquots were transferred into glass tubes and the efflux pump inhibitor carbonyl cyanide 3-chlorophenylhydrazone (CCCP) was added at a final concentration of 25 mM (5 mM stock solution in 50% DMSO). After 15 min 5 mM Nile red dye was added (a stock solution of 5 mM in 10 % dimethyl formamide, 90 % ethanol) and the cell suspension was incubated on a shaker (140 r. p. m.; 30 °C) for 3 h, followed by a 60 min incubation without shaking at room temperature and centrifugation for 5 min at 4,400 r. p. m. in the tabletop centrifuge. Supernatant was entirely removed and cells were resuspended in 1 ml PPB-Mg (or in PPB-Mg supplemented with increased Tc concentrations, see Fig. EV2). Immediately thereafter 0.3 ml of this cell suspension was transferred to a 96 well microtiter plate and 15 μl of 1 M glucose, was added to trigger Nile red efflux. Fluoresence of the cell suspension was followed over 1500 sec (excitation at 552 n., and emission at 636 nm) on the Tecan reader. Three independent experiments revealed similar results..

### Analysis of the antisense promoter P_as_

Plasmid pSUP-PasRegfp containing the transcriptional fusion of *egfp* to promoter P_as_ was used to analyze the inducubility of the promoter by antimicrobial compounds, which were added to cultures at the OD_600 nm_ of 0.5. Since this plasmid confers Tc resistance, it was necessary to incubate strain 2011 (pSUP-PasRegfp) with the chromosomally integrated plasmid overnight without Tc (essentially all cells retained the plasmid, as confirmed by qPCR analysis), before Tc was added at the indicated concentrations. Similarly, other antimicrobial compounds were added at subinhibitory concentrations (see above) to 2011 (pSUP-PasRegfp) cultures that were incubated without Tc overnight. RNA was isolated before (time point 0) and at the indicated time points after antibiotic addition, and changes in the reporter *egfp* mRNA upon exposure was analyzed by qRT-PCR.

### RNA purification

For Northern blot hybridization and qRT-PCR analysis, total RNA of *S. meliloti* and *A. tumefaciens* was purified from 15 ml culture (OD_600_ = 0.5). The cells were cooled by adding the culture directly into tubes with ice rocks (corresponding to a volume of 15 ml). After a centrifugation at 6,000 g for 10 min at 4 °C, the pellet was resuspended in 250 μl TRIzol (Life Technologies, Darmstadt, Germany). Lysis was performed with in a Laboratory Mixer Mill (Retsch MM200) (4 °C) with glass beads, two times for 15 min, interrupted by incubation at 65 °C for 10 min. Then 750 μl TRIzol was added to the samples and RNA was isolated according to the manufacturer’s instructions. Residual RNases were removed by additional extraction with hot-phenol, phenol:chloroform:isoamylalcohol (25:24:1) and chloroform:isoamylalcohol (24:1). RNA was ethanol-precipitated and dissolved in ultrapure water. For RNA half-live measurements, 1 ml *S. meliloti* or *A. tumefaciens* culture was added to 2 ml RNAprotect Bacteria Reagent (Qiagen) and RNA was isolated using RNeasy columns (Qiagen). RNA from *B. japonicum* was isolated with hot phenol [62]. For analysis of the concentration and purity of RNA, absorbance at 260 nm and 280 nm was measured. Integrity of the isolated RNA was controlled by separation in 10% polyacrylamide-urea gels and staining with ethidium bromide. For qRT-PCR analysis, residual DNA was removed by incubation of 10 μg RNA with 1 μl TURBO-DNase (Ambion) for 30 minutes. Prior to the qRT-PCR analysis, the RNA samples were tested for presence of DNA by PCR with *rpoB*-specific primers. The DNA-free RNA was then diluted to a concentration of 20 ng/μl and stored at −20°C.

### Northern Blot hybridization

Total RNA (10 μg per lane) was separated in 10% polyacrylamide-urea gels, transferred to a positively charged nylon membrane and cross-linked by UV. After prehybridization for 2 h at 56 °C with a buffer containing 6× SSC, 2.5× Denhardt’s solution, 1% SDS and 10 μg/ml salmon sperm DNA, hybridization was performed with radioactively labeled oligonucleotides (Table S2) in a solution containing 6× SSC, 1% SDS, 10 μg/ml salmon sperm DNA for at least 6 h at 56 °C. Membrane washing was performed twice for 2 to 5 min in 0.01% SDS, 5× SSC at room temperature. Signals were detected using a BioRad molecular imager. For rehybridization, membranes were washed in 0.1% SDS for 20 min at 96°C.

### Radioactive labeling of oligonucleotide probes

Oligonucleotide (10 pmol) were labeled at the 5′-terminus using 15 μCi [γ-^32^P]-ATP (Hartmann Analytics, Braunschweig, Germany) and 5 U T4 polynucleotide kinase in a 10 μl reaction mixture, which was incubated for 60 min at 37 °C. After adding 30 μl water, unincorporated nucleotides were removed using MicroSpin G-25 columns (GE Healthcare Life Sciences).

### *In vitro* transcription

For *in vitro* transcription the MEGAshortscript T7 kit (Thermo Fisher Scientific, Vilnius, Lithuania) was used. The T7 promoter sequence was integrated in one of the primers for PCR amplification of the template (see Appendix Table S2), which was column-purified and eluted in ultrapure water. The transcription reaction contained 500 ng template, 1x T7-polymerase buffer, 7.5 mM ATP, 7.5 mM CTP, 7.5 mM GTP, 7.5 mM UTP and 25 U T7-enzyme mix. It was incubated for at least 5 h at 37°C. To remove the DNA template, 1 μl TURBO-DNase was added and the mixture was incubated for 1h at 37°C. The *in vitro* transcript was extracted with acidic phenol, precipitated with ethanol and analyzed in a 10% polyacrylamide-urea gel stained with ethidium bromide.

### Strand-specific, real time, reverse transcriptase PCR (qRT-PCR)

Relative steady-state levels of specific RNAs by real-time RT-PCR (qRT-PCR) were analyzed using the Brilliant III Ultra Fast SYBR® Green QRT-PCR Mastermix (Agilent, Waldbronn, Germany). Strand-specific analysis was performed as follows: 5 μl Master Mix (supplied), 0.1 μl DTT (100 mM; supplied), 0.5 μl Ribo-Block solution (supplied), 0.4 μl water, 1 μl of the reverse primer (10 pmol/μl), and 2 μl RNA (20 ng/μl) were assembled in a 9-μl reaction mixture. After cDNA synthesis the reverse transcriptase was inactivated by incubation for 10 min at 96 °C. Then the samples were cooled to 4°C, 1 μl of the second primer (10 pmol) was added, and real-time PCR was performed starting with 5 min incubation at 96 °C. The efficiencies of the used primer pairs (Appendix Table S2) were determined by PCR of serial two-fold RNA dilutions. Primer pairs were designed using Primer3 [63]. The qRT-PCR reactions were conducted in a spectrofluorometric thermal cycler (BioRad, München, Germany). The quantification cycle (Cq) was set to a cycle at which the curvature of the amplification is maximal [64]. For determination of steady-state mRNA levels, *rpoB* (encodes the β subunit of RNA polymerase) was used as a reference gene [56]. For half-life determination, the stable, but highly abundant 16S rRNA was used as a reference molecule. Therefore (to achieve similar Cq of mRNA and 16S rRNA), the 10-μl reaction for qRT-PCR with 16S rRNA specific primers contained 2 μl RNA with a concentration of 0.002 ng/μl [39]. The Pfaffl formula was used in to calculate fold changes of mRNA amounts [65]. The qRT-PCRs reactions with an RNA sample were performed in technical replicates. If the Cq difference between the technical replicates was > 0.5, the analysis was repeated. In such a case, the RNA sample of the outliers and, as a control, at least one of the other RNA samples, were analyzed once again by qRT-PCR. If the Cq difference of the reference gene in independent biological experiments was > 1 (for *rpoB*) or > 2 (for 16S rRNA), the analysis was repeated. qPCR product specificity was validated by a melting curve after the qPCR-reaction and by gel electrophoresis. No-template controls and negative mRNA controls (RNAs expected to be not affected under the applied conditions, e.g. *trpDC* mRNA which is transcribed from a second *trp* operon and is regulated by rnTrpL but not by peTrpL [39]) were always included.

For analysis of total RNA, the qRT-PCR of the gene of interest (e.g. *smeR*) and of the reference gene *rpoB* were performed using portions of the same DNA-free RNA sample, and log_2_fold changes of mRNA levels after induction by IPTG and/or exposure to antibiotics were determined. For analysis of coimmunoprecipitated RNA, the qRT-PCR of the gene of interest was performed using a CoIP-RNA sample, while total RNA of the same culture (harvested prior to cell lysis for CoIP) was used for the *rpoB* qRT-PCR. Then, the Pfaffl formula was used to calculate the fold enrichment of specific RNAs by CoIP with 3×FLAG-peTrpL, in comparison to the corresponding mock CoIP.

### mRNA half-life determination

Stability of mRNA was determined as described [39]. Rifampicin was added to a final concentration of 800 μg/ml (stock concentration 150 mg/ml in methanol) to stop cellular transcription. Culture aliquots were withdrawn at time points 0, 2, 4 and 6 min and RNA was isolated. To determine the relative levels of specific mRNAs, qRT-PCR analysis with 16S rRNA as a reference was performed (see above). Linear-log graphs were used for half-lives calculation.

### Real time PCR (qPCR)

Plasmid-specific primers (Appendix Table S2) were used to test whether the chromosomally integrated plasmid pSUP-PasRegfp is lost after culture incubation without selective pressure overnight. As a reference gene *rpoB* was used. Power SYBR® PCR Mastermix (Qiagen) was used for the qPCR reactions. The template and primer concentrations, reaction conditions and quantification were performed as described for qRT-PCR of total RNA.

### Coimmunoprecipitation of RNA with 3×FLAG-peTrpL and its analysis

The CoIP of RNA that was used for RNA-seq analysis was performed with the two-plasmid strain 2011 (pSRKGm-3×FLAG-peTrpL, pRK4352) which was cultivated in medium with Gm and Tc (20 μg/ml). Cells were harvested 10 min after induction of 3×FLAG-peTrpL production with IPTG. In parallel, strain 2011 (pSRKGm-peTrpL, pRK4352) was cultivated and treated similarly for the mock CoIP control. Cell pellets were resuspended in 5 ml buffer A (20 mM Tris, pH 7.5, 150 mM KCl, 1 mM MgCl_2_, 1 mM DTT) containing 10 mg/ml lysozyme, 2 μg/ml Tc and 1 tablet of protease inhibitor cocktail (Sigma-Aldrich, St. Louis, MO, USA) per 40 ml buffer. After lysis by sonication, 40 μl Anti FLAG® M2 Magnetic Beads (Sigma-Aldrich, Cat#SLBT7133) were added to the cleared lysate and incubated for 2 h at 4 °C. Then the beads were split into two portions: one of them was washed 3 times with 500 μl buffer A containing 2 μg/ml Tc, while the other was washed with buffer without Tc. Protease inhibitors were included in the first two washing steps. Finally, the beads were resuspended in 50 μl buffer A. Coimmunoprecipitated RNA (CoIP-RNA) was purified using TRIzol, without subsequent hot-phenol treatment. The CoIP-RNA absorbance at 260 and 280 nm suggested that efficient coimmunoprecipitation of RNA was possible only when Tc was present in the washing buffer. Therefore, only CoIP-RNA from beads washed with Tc-containing buffer was subjected to RNA-seq analysis (Vertis Biotechnologie AG, Freising, Germany). cDNA reads were mapped as described [66]. The CoIP-RNA from beads washed with or without Tc in the buffer was compared by qRT-PCR (see above).

Strain 2011 (pSRKGm-3×FLAG-peTrpL) and the corresponding mock control 2011 (pSRKGm - peTrpL), which were cultivated in medium with Gm only, were also used for FLAG-CoIP. In this case, for successful CoIP, subinhibitory concentrations of the used antimicrobial compounds (see above) were added to the cultures along with IPTG, and to the washing buffer of the CoIP procedure. CoIP-RNA was analyzed by qRT-PCR.

### ARNP complex reconstitution

Used peTrpL and 3×FLAG-peTrpL peptides were synthesized by Thermo Fischer Scientific (Darmstadt, Germany). 10 mg peTrpL was dissolved in 50 μl acetonitrile and 950 μl ultrapure water was added. 1 mg 3×FLAG-peTrpL was dissolved in 1 ml 50 % DMSO. 50 μl-aliquots were stored at − 20 °C. Peptides were diluted in ultrapure water prior to usage. For reconstitution, 100 ng mini-*smeR in vitro* transcript (4.4 pmol), 100 ng mini-As-smeR *in vitro* transcript (4.4 pmol), 50 ng peTrpL (27 pmol) and 50 ng 3×FLAG-peTrpL (11 pmol) were mixed in buffer B (20 mM Tris, pH 8.0, 150 mM KCl, 1 mM MgCl_2_, 1 mM DTT), in a volume of 48 μl. Then 2 μl antibiotic or flavonoid solution was added. To negative control samples, 2 μl ethanol, methanol or water (the solvents of the antibiotic solutions) were added. Following final concentrations of the antimicrobial compounds were used: 1.5 μg/ml Tc, 27 μg/ml Em, 9 μg/ml Cl, 45 μg/ml Km, 90 μg/ml Gs, and 45 μg/ml Lt. The samples were incubated for 20 min at 20 °C, and then 3×FLAG-peTrpL-containing complexes were isolated by CoIP with anti-FLAG antibodies. The antimicrobial compounds were present in the washing buffer in the concentrations given above. After extensive washing, RNA was purified and analyzed by Northern blot hybridization.

### Analysis of the conservation of peTrpL function

Phyre^2^ [67] was used to analyze SmeR, leading to its identification as a TtgR [18] homolog. Closest homologs of the *S. meliloti smeR* gene in *A. tumefaciens* and *B. japonicum* were identified by Blastp (https://blast.ncbi.nlm.nih.gov/Blast.cgi). In *A. tumefaciens*, this was the gene Atu3201 (*acrR*), which encodes a TetR-type repressor and is a part of the *acrABR* operon additionally encoding a MDR efflux pump [13]. In *B. japonicum*, the best match was the orphan gene blr2396 encoding a TetR-type repressor, and thus blr2396. was analyzed as a *smeR* homolog. Plasmid pSRKTc-Atu-peTrpL was used to induce by IPTG the Atu-peTrpL production in *A. tumefaciens* for 10 min. Changes in mRNA levels were calculated in comparison to 0 min. Due to the lack of a suitable inducible system for *B. japonicum*, Bja-peTrpL was overproduced constitutively from the chromosomally integrated, Tc-resistance-conferring plasmid pRJ-Bja-rnTrpL. Changes in mRNA levels were calculated in comparison to the EVC. Phenotypic changes were tested as indicated.

## Data availability

The RIP-Seq data discussed in this publication have been deposited in NCBI's Gene Expression Omnibus [68]; accession number GSE118689.

## Acknowledgments

We are grateful to Janina Gerber and Maximilian Stötzel for help in some experiments. This work was funded by DFG (Ev 42/6-1; Ev 42/7-1 in SPP2002; GRK2355 project number 325443116). S.L. was supported by the China Scholarship Council (No. 201708080082)

## Author contributions

Conceptualization, E.EH., H.M.; Methodology, E.E.H., H.M.; Investigation, H.M., S.A., S.B.W., S.L.; Data curation, K.U.F., Formal analysis, H.M., K.U.F., S.L.; Writing E.E.H., H.M; Visualization, E.E.H., H.M., S.L.; Supervision, funding acquisition and project administration, E.E.H.

## Conflict of interests

The authors declare no conflict of interests.

## Expanded View Figure legends

**Figure EV1.**
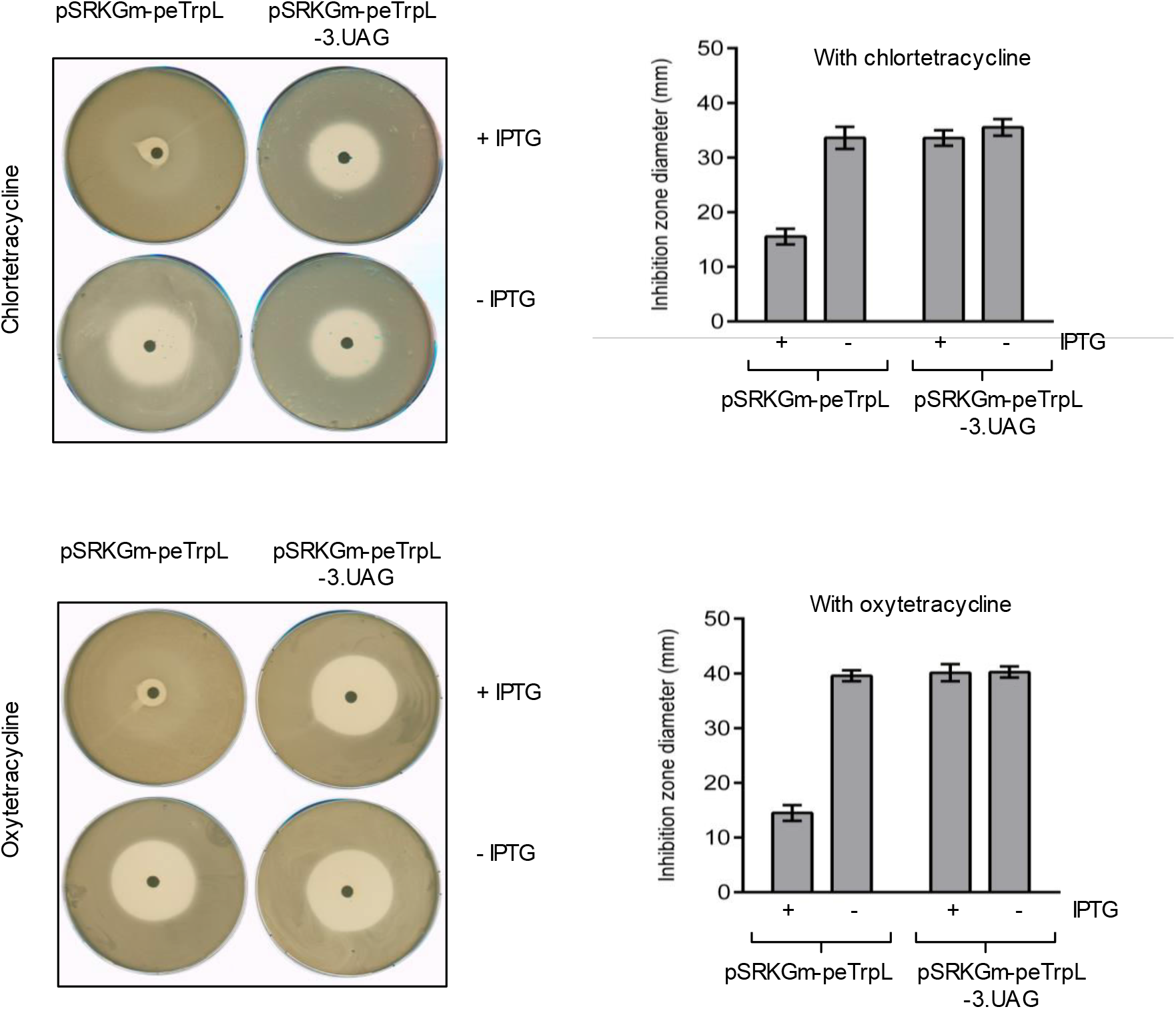
Increased resistance to natural tetracyclines upon induced peTrpL overproduction. Representative plates with zones of growth inhibition caused by the centrally applied chlortetracycline and oxytetracycline are shown. Strain *S. meliloti* 2011*ΔtrpL* containing the indicated plasmids was used. Presence of IPTG in the growth media is indicated. On the right side, data from three independent cultures are presented as mean ± s.d.

**FigureEV2.**
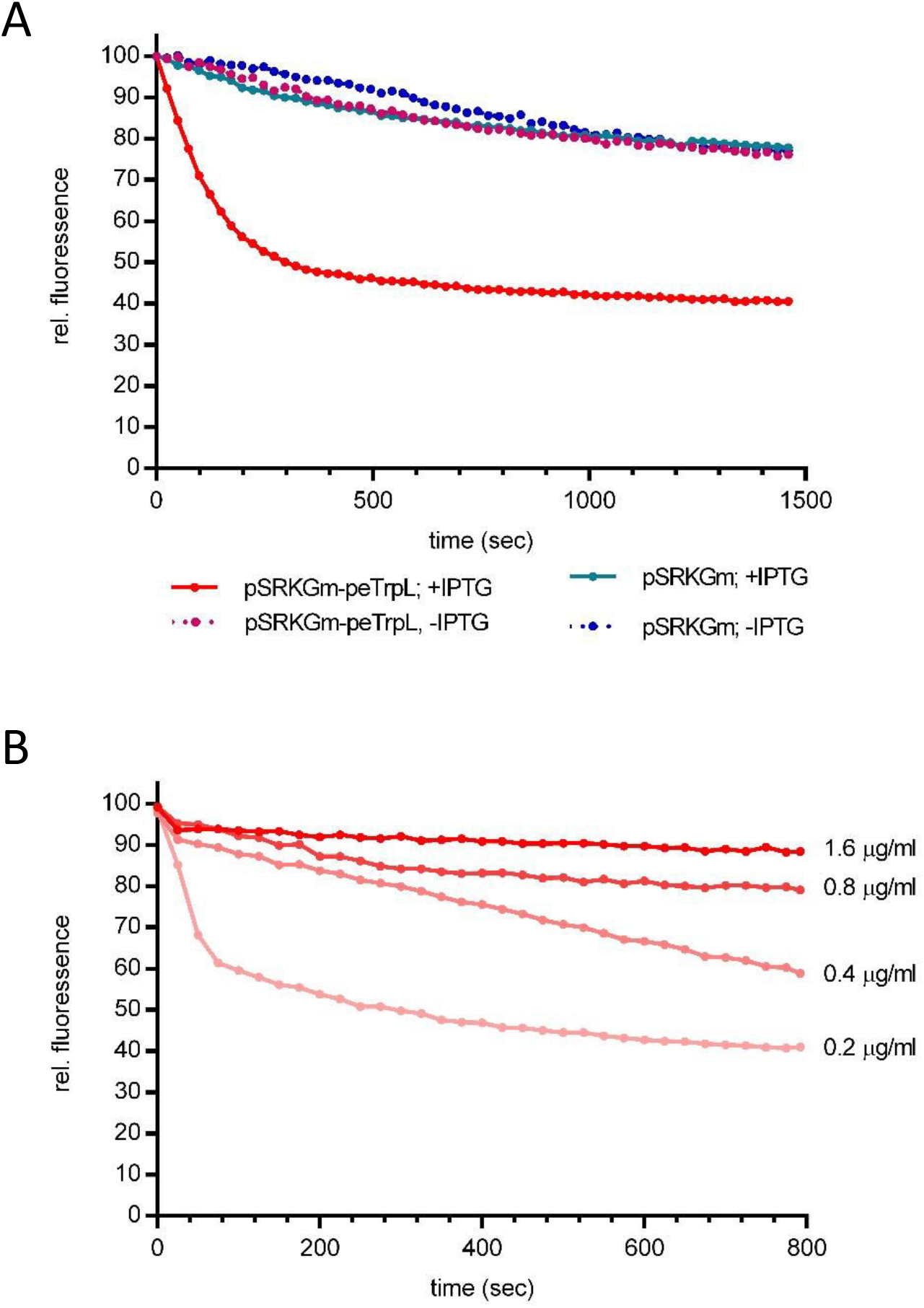
Induced peTrpL production increases the cellular efflux. A Nile red efflux assay for determination of the efflux activity of strain 2011 (pSRKGm-peTrpL, pRK4352) which was grown in the presence or absence of IPTG. pSRKGm, EVC. The Nile red dye generates a fluorescence signal only if present in the cell. Fast decline of fluorescence over time correlates with high efflux pump activity. Data of a representative experiment is shown. B Tc competes with Nile red for the efflux, showing that they are extruded by the same pump(s). Nile red efflux assay was performed with strain 2011 (pSRKGm-peTrpL, pRK4352) cultivated with IPTG. Increasing Tc concentrations were added together with glucose to start the efflux.

**Figure EV3.**
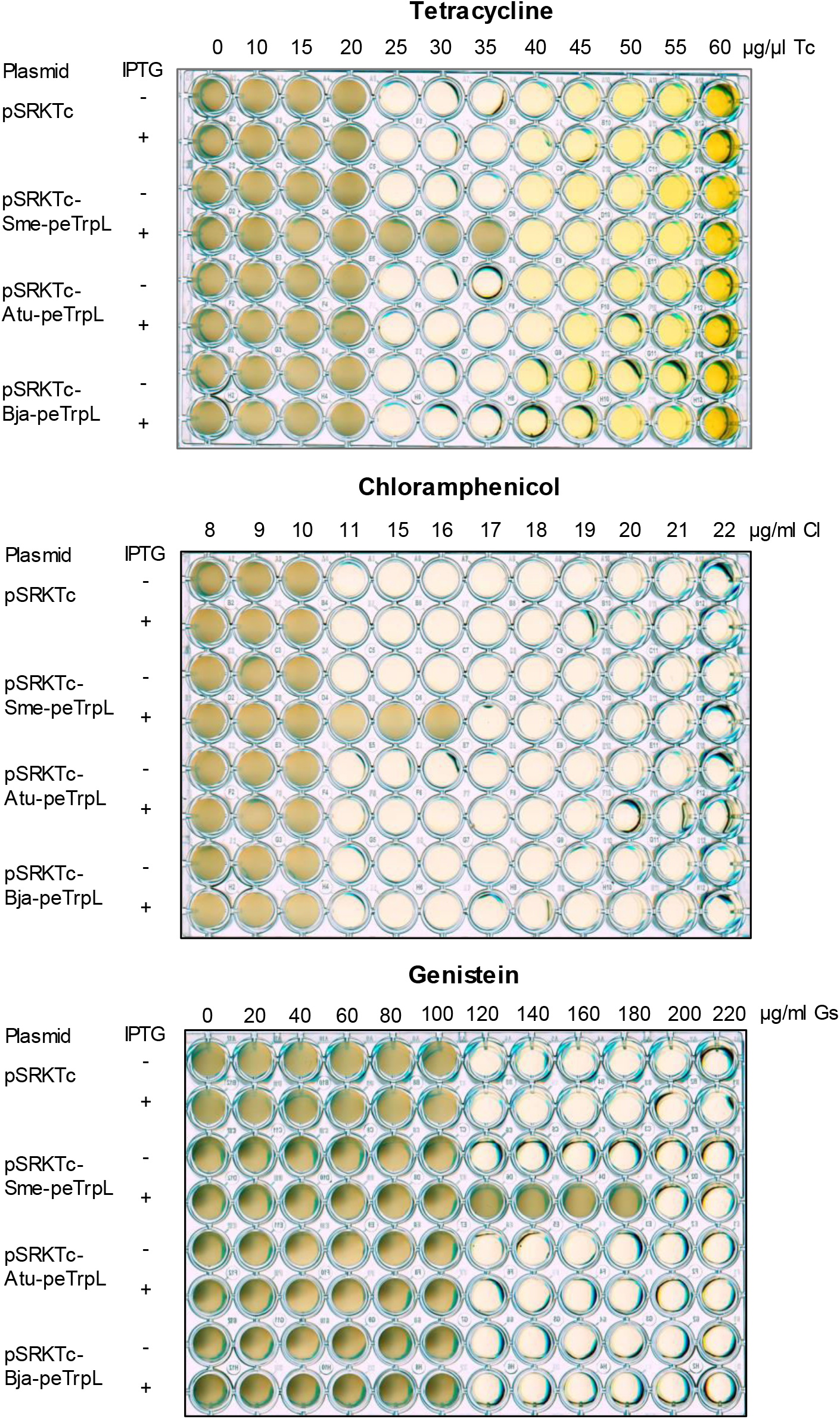
Heterologous peTrpL peptides do not increase the multiresistance of *S. meliloti*. Shown are representative plates with strain 2011*ΔtrpL* containing the indicated plasmids. Used antimicrobial compounds, their concentrations and presence of IPTG in the medium are indicated.

## References

1. Hutchings M, Truman A, Wilkinson B (2019) Antibiotics: past, present and future. Curr Opin Microbiol 13: 72–80

2. Genilloud O (2019) Natural products discovery and potential for new antibiotics. Curr Opin Microbiol 15: 81–87

3. Christaki E, Marcou M, Tofarides A (2019) Antimicrobial Resistance in Bacteria: Mechanisms, Evolution, and Persistence. J Mol Evol 2019 Oct 28

4. Blair JM, Webber MA, Baylay AJ, Ogbolu DO, Piddock LJ (2015) Molecular mechanisms of antibiotic resistance. Nat Rev Microbiol 13: 42–51

5. Peterson E, Kaur P (2018) Antibiotic Resistance Mechanisms in Bacteria: Relationships Between Resistance Determinants of Antibiotic Producers, Environmental Bacteria, and Clinical Pathogens. Front Microbiol 9: 2928

6. Paulsen IT (2003) Multidrug efflux pumps and resistance: regulation and evolution. Curr Opin Microbiol 6: 446–451

7. Piddock LJ (2006) Multidrug-resistance efflux pumps - not just for resistance. Nat Rev Microbiol 4: 629–636

8. Murakami, S., Nakashima, R., Yamashita, E., Matsumoto, T. & Yamaguchi, A. Crystal structures of a multidrug transporter reveal a functionally rotating mechanism. Nature 443, 173–179 (2006)

9. Nakashima, R., Sakurai, K., Yamasaki, S., Nishino, K. & Yamaguchi, A. Structures of the multidrug exporter AcrB reveal a proximal multisite drug-binding pocket. Nature 480, 565–569 (2011).

10. Eicher, T. et al. Transport of drugs by the multidrug transporter AcrB involves an access and a deep binding pocket that are separated by a switch-loop. Proc. Natl Acad. Sci. USA 109, 5687–5692 (2012).

11. Walsh F, Duffy B (2013) The culturable soil antibiotic resistome: a community of multi-drug resistant bacteria. PLoS One 8: e65567

12. Naamala J, Jaiswal SK, Dakora FD (2016) Antibiotics Resistance in Rhizobium: Type, Process, Mechanism and Benefit for Agriculture. Curr Microbiol. 72: 804–816

13. Nuonming P, Khemthong S, Dokpikul T, Sukchawalit R, Mongkolsuk S (2018) Characterization and regulation of AcrABR, a RND-type multidrug efflux system, in Agrobacterium tumefaciens C58. Microbiol Res 214: 146–155

14. Eda S, Mitsui H, Minamisawa K (2011)Involvement of the smeAB multidrug efflux pump in resistance to plant antimicrobials and contribution to nodulation competitiveness in Sinorhizobium meliloti. Appl Environ Microbiol. 77: 2855–2862

15. Hinrichs W, Kisker C, Düvel M, Müller A, Tovar K, Hillen W, Saenger W (1994) Structure of the Tet repressor-tetracycline complex and regulation of antibiotic resistance. Science 264: 418–420

16. Terán W, Felipe A, Segura A, Rojas A, Ramos JL, Gallegos MT (2003) Antibiotic-dependent induction of Pseudomonas putida DOT-T1E TtgABC efflux pump is mediated by the drug binding repressor TtgR. Antimicrob Agents Chemother. 47: 3067–3072.

17. Lin J, Akiba M, Sahin O, Zhang Q (2005) CmeR functions as a transcriptional repressor for the multidrug efflux pump CmeABC in Campylobacter jejuni. Antimicrob Agents Chemother 49: 1067–1075

18. Alguel Y, Meng C, Terán W, Krell T, Ramos JL, Gallegos MT, Zhang X. (2007) Crystal structures of multidrug binding protein TtgR in complex with antibiotics and plant antimicrobials. J Mol Biol 369: 829–840

19. Alekshun MN, Levy SB (1997) Regulation of chromosomally mediated multiple antibiotic resistance: the mar regulon. Antimicrob Agents Chemother. 41: 2067–2075

20. Kwak JH, Choi EC, Weisblum B (1991) Transcriptional attenuation control of ermK, a macrolide-lincosamide-streptogramin B resistance determinant from Bacillus licheniformis. J Bacterio. 173: 4725–4735

21. Schwarz S, Kehrenberg C, Doublet B, Cloeckaert A (2004) Molecular basis of bacterial resistance to chloramphenicol and florfenicol. FEMS Microbiol Rev 28: 519–542

22. Dar D, Shamir M, Mellin JR, Koutero M, Stern-Ginossar N, Cossart P, et al (2016) Term-seq reveals abundant ribo-regulation of antibiotics resistance in bacteria. Science 352: aad9822

23. Dersch P, Khan MA, Mühlen S, Görke B (2017) Roles of Regulatory RNAs for Antibiotic Resistance in Bacteria and Their Potential Value as Novel Drug Targets. Front Microbiol 8: 803

24. Khan MA, Göpel Y, Milewski S, Görke B (2016) Two small RNAs conserved in Enterobacteriaceae provide intrinsic resistance to antibiotics targeting the cell wall biosynthesis enzyme glucosamine-6-phosphate synthase. Front Microbiol 7: 908

25. Kim T, Bak G, Lee J, Kim KS (2015) Systematic analysis of the role of bacterial Hfq-interacting sRNAs in the response to antibiotics. J Antimicrob Chemother 70: 1659–1668

26. Nishino K, Yamasaki S, Hayashi-Nishino M, Yamaguchi A (2011) Effect of overexpression of small non-coding DsrA RNA on multidrug efflux in Escherichia coli. J Antimicrob Chemother 66: 291– 296

27. Yamada J, Yamasaki S, Hirakawa H, Hayashi-Nishino M, Yamaguchi A, and Nishino K (2010) Impact of the RNA chaperone Hfq on multidrug resistance in Escherichia coli. J Antimicro. Chemother 65: 853–858

28. El-Mowafi SA, Alumasa J N, Ades SE, Keiler KC (2014) Cell-based assay to identify inhibitors of the Hfq-sRNA regulatory pathway. Antimicrob Agents Chemother 58: 5500–5509

29. Yu J, Schneiders T (2012) Tigecycline challenge triggers sRNA production in Salmonella enterica serovar Typhimurium. BMC Microbiol 12: 195

30. Howden BP, Beaume M, Harrison PF, Hernandez D, Schrenzel J, Seemann T, Francois P, Stinear TP (2013) Analysis of the small RNA transcriptional response in multidrug-resistant Staphylococcus aureus after antimicrobial exposure. Antimicrob Agents Chemother 57: 3864–3874

31. Orr MW, Mao Y, Storz G, Qian SB (2019) Alternative ORFs and small ORFs: shedding light on the dark proteome. Nucleic Acids Res pii: gkz734

32. Storz G, Wolf YI, Ramamurthi KS (2014) Small proteins can no longer be ignored. Annu Rev Biochem 83: 753–777

33. Levin PA, Fan N, Ricca E, Driks A, Losick R, Cutting S (1993) An unusually small gene required for sporulation by Bacillus subtilis. Mol Microbiol 9: 761–771

34. Hobbs EC, Yin X, Paul BJ, Astarita JL, Storz G (2012) Conserved small protein associates with the multidrug efflux pump AcrB and differentially affects antibiotic resistance. Proc Natl Acad Sci U S A 109: 16696–16701

35. Vitreschak AG, Lyubetskaya EV, Shirshin MA, Gelfand MS, Lyubetsky VA (2004) Attenuation regulation of amino acid biosynthetic operons in proteobacteria: comparative genomics analysis. FEMS Microbiol Lett 234: 357–370

36. Merino E, Jensen RA, Yanofsky C (2008) Evolution of bacterial trp operons and their regulation. Curr Opin Microbiol 11: 78–86

37. Yanofsky C (1981) Attenuation in the control of expression of bacterial operons. Nature 289: 751–758

38. Bae YM, Crawford IP (1990) The Rhizobium meliloti trpE(G) gene is regulated by attenuation, and its product, anthranilate synthase, is regulated by feedback inhibition. J Bacteriol 172: 3318–3327

39. Melior H, Li S, Madhugiri R, Stötzel M, Azarderakhsh S, Barth-Weber S, Baumgardt K, Ziebuhr J, Evguenieva-Hackenberg E (2019) Transcription attenuation-derived small RNA rnTrpL regulates tryptophan biosynthesis gene expression in trans. Nucleic Acids Res 47: 6396–6410

40. Nguyen F, Starosta AL, Arenz S, Sohmen D, Dönhöfer A, Wilson DN (2014) Tetracycline antibiotics and resistance mechanisms. Biol Chem 395: 559–575

41. Zahn, K. (1996) Overexpression of an mRNA dependent on rare codons inhibits protein synthesis and cell growth. J Bacteriol 178, 2926–2933

42. Terán W, Krell T, Ramos JL, Gallegos MT (2006) Effector-repressor interactions, binding of a single effector molecule to the operator-bound TtgR homodimer mediates derepression. J Biol Chem 281:7102–7109

43. Schlüter JP, Reinkensmeier J, Barnett MJ, Lang C, Krol E, Giegerich R, Long SR, Becker A (2013) Global mapping of transcription start sites and promoter motifs in the symbiotic α-proteobacterium Sinorhizobium meliloti BMC Genomics 14: 156

44. Casse F, Boucher C, Julliot JS, Michel M, Denarie J (1979) Identification and characterization of large plasmids in Rhizobium meliloti using agarose-gel electrophoresis. Gen Microbiol 113: 229–242

45. Young JM (2003) The genus name Ensifer Casida 1982 takes priority over Sinorhizobium Chen et al. 1988, and Sinorhizobium morelense Wang et al. 2002 is a later synonym of Ensifer adhaerens Casida 1982. Is the combination “Sinorhizobium adhaerens” (Casida 1982) Willems et al. 2003 legitimate? Request for an Opinion. Int J Syst Evol Microbiol 53: 2107–2110

46. Cha C, Gao P, Chen YC, Shaw PD, Farrand SK (1998) Production of acyl-homoserine lactone quorum-sensing signals by gram-negative plantassociated bacteria. Mol Plant Microbe Interact 11: 1119–1129

47. Lassalle F, Campillo T, Vial L, Baude J, Costechareyre D, Chapulliot D, Shams M, Abrouk D, Lavire C, Oger-Desfeux C, et al (2011) Genomic species are ecological species as revealed by comparative genomics in Agrobacterium tumefaciens. Genome Biol Evol 3: 762–781

48. Beringer JE (1974) R factor transfer in Rhizobium leguminosarum. J Gen Microbiol 84: 188–198

49. Regensburger B, Hennecke H (1983) RNA polymerase from Rhizobium japonicum. Arch Microbiol 135: 103–109

50. Delamuta JR, Ribeiro RA, Ormeño-Orrillo E, Melo IS, Martínez-Romero E, Hungria M (2013) Polyphasic evidence supporting the reclassification of Bradyrhizobium japonicum group Ia strains as Bradyrhizobium diazoefficiens sp. nov. Int J Syst Evol Microbiol 63: 3342–3351

51. Mesa S, Hauser F, Friberg M, Malaguti E, Fischer HM, Hennecke H (2008) Comprehensive assessment of the regulons controlled by the FixLJ-FixK_2_-FixK_1_ cascade in Bradyrhizobium japonicum. J Bacteriol 190: 6568–6579

52. Sambrook J, Fritsch EF, Maniatis T (1989) Molecular cloning: A laboratory manual. 2. Cold Spring Harbor Laboratory Press, Cold Spring Harbor, NY

53. Khan SR, Gaines J, Roop RM2^nd^, Farrand SK (2008) Broad-host-range expression vectors with tightly regulated promoters and their use to examine the influence of TraR and TraM expression on Ti plasmid quorum sensing. Appl Environ Microbiol 74: 5053–5062

54. Fischer HM, Babst M, Kaspar T, Acuna G, Arigoni F, Hennecke H(1993) One member of a gro-ESL-like chaperonin multigene family in Bradyrhizobium japonicum is co-regulated with symbiotic nitrogen fixation genes. EMBO J 12: 2901–2912

55. Mank NN, Berghoff BA, Hermanns YN, Klug G (2012) Regulation of bacterial photosynthesis genes by the small noncoding RNA PcrZ. Proc. Natl Acad Sci U S A 109:16306–16311

56. Baumgardt K, Šmídová K, Rahn H, Lochnit G, Robledo M, Evguenieva-Hackenberg E (2016) The stress-related, rhizobial small RNA RcsR1 destabilizes the autoinducer synthase encoding mRNA sinI in Sinorhizobium meliloti. RNA Biol 13:486–499

57. Hahn J, Thalmann S, Migur A, von Boeselager RF, Kubatova N, Kubareva E, Schwalbe H, Evguenieva-Hackenberg E (2017) Conserved small mRNA with an unique, extended Shine-Dalgarno sequence. RNA Biol 14: 1353–1363

58. McIntosh M, Krol E, Becker A (2008) Competitive and cooperative effects in quorum-sensing-regulated galactoglucan biosynthesis in Sinorhizobium meliloti. J Bacteriol 190: 5308–5317

59. Schlüter JP, Czuppon P, Schauer O, Pfaffelhuber P, McIntosh M, Becker A (2015) Classification of phenotypic subpopulations in isogenic bacterial cultures by triple promoter probing at single cell level. J Biotechnol 198: 3–14

60. Simon R, Priefer U, Pühler A (1982) A broad host range mobilization system for in vivo genetic engineering: transposon mutagenesis in gram-negative bacteria. Biotechnology 1: 784–791

61. Bohnert JA, Karamian B, Nikaido H (2010) Optimized Nile Red Efflux Assay of AcrAB-TolC Multidrug Efflux System Shows Competition between substrates. Antimicrob Agents Chemother 54: 37770–3775

62. von Gabain A, Belasco JG, Schottel JL, Chang AC, Cohen SN (1983) Decay of mRNA in Escherichia coli: investigation of the fate of specific segments of transcripts. Proc Natl Acad Sci U S A 80: 653–657

63. Untergasser A, Nijveen H, Rao X, Bisseling T, Geurts R, Leunissen JA (2007) Primer3Plus, an enhanced web interface to Primer3. Nucleic Acids Res 35:W71–74

64. Bustin SA, Benes V, Garson JA, Hellemans J, Huggett J, Kubista M, Mueller R, Nolan T, Pfaffl MW, Shipley GL et al (2009) The MIQE guidelines: minimum information for publication of quantitative real-time PCR experiments. Clin. Chem. 55: 611–622

65. Pfaffl MW(2001) A new mathematical model for relative quantification in real-time RT-PCR. Nucleic Acids Res. 29: e45

66. Sharma CM, Hoffmann S, Darfeuille F, Reignier J, Findeiss S, Sittka A, Chabas S, Reiche K, Hackermüller J, Reinhardt R, et al (2010) The primary transcriptome of the major human pathogen Helicobacter pylori. Nature 464: 250–255

67. Kelley LA, Mezulis S, Yates CM, Wass MN, Sternberg MJ (2015) The Phyre2 web portal for protein modeling, prediction and analysis. Nat Protoc 10: 845–858

68. Edgar R, Domrachev M, Lash AE (2002) Gene Expression Omnibus: NCBI gene expression and hybridization array data repository. Nucleic Acids Res 30: 207–210

